# CD27⁺ γδ T Cells Drive Plaque Instability in Advanced Atherosclerosis: Targeting CXCR3 for Therapeutic Intervention

**DOI:** 10.1101/2025.08.19.671168

**Authors:** Peter Kanellakis, Anh Cao, Yi Ee Lye, Ban-Hock Toh, Alex Bobik, Tin Kyaw

## Abstract

**Background:** Atherosclerosis is a chronic inflammatory disease of the arterial wall that underlies most myocardial Ischaemic events. While multiple immune subsets contribute to plaque progression and instability, the role of γδ T cells remains poorly understood. We examined the contribution of γδ T cells to lesion development, progression and instability, and explored the therapeutic potential of their pharmacological blockade.

**Methods:** To investigate the role of γδ T cells, chimeric atherosclerosis-prone mice lacking γδ T cells were utilized in both loss- and gain-of-function experiments. Mixed bone marrow chimeras were generated to assess the role of γδ T cell-derived interferon-γ (IFN-γ) and perforin (Pfp). The therapeutic efficacy of AMG487 on plaque stability was evaluated in a preclinical tandem-stenosis mouse model. Lesion size, plaque composition, and stability were assessed using histology, immunoassays, and molecular biology techniques.

**Results:** CD27+ γδ T cells accumulated in atherosclerotic lesions and promoted plaque progression and instability via IFN-γ- and Pfp-dependent manners. As early infiltrators, they amplified necrosis and inflammation by enhancing immune cell recruitment, thereby exacerbating lesion vulnerability. CXCR3 antagonism with AMG487 inhibited γδ T cell recruitment to plaques, reduced lesion size, and promoted features of plaque stability, including increased smooth muscle cell content and thicker fibrous caps.

**Conclusions:** CD27⁺ γδ T cells, which promote inflammation and necrosis through both direct and indirect mechanisms, are key drivers of plaque progression and instability. Targeting their recruitment via CXCR3 blockade enhances plaque stability and may represent a promising therapeutic strategy to reduce the risk of myocardial infarction.

## Introduction

Myocardial infarction (MI) is the leading cause of global mortality^1,2^, and results from acute coronary ischemia typically following sudden occlusion of atherosclerotic coronary arteries. While lowering low-density lipoprotein (LDL) cholesterol through diet and therapeutic agents is the primary strategy for preventing MI in high-risk patients, a substantial fraction of patients still suffer Ischaemic events despite achieving target lipid levels^3,4^. This residual risk highlights the role of non-lipid factors, most notably inflammation and immune mechanisms, in driving plaque destabilisation and rupture^5,6^. Our group and others have linked diverse immune cell subsets to chronic arterial inflammation and plaque vulnerability^7–14^, underscoring the importance of targeting maladaptive immune responses to prevent and treat MI.

γδ T cells, though a minor fraction of the T-cell pool, possess both innate-like rapid responses and potent adaptive specificity. They respond directly to peptide antigens independent of major histocompatibility complex (MHC) molecules^15^, yet can also recognise peptide antigens via MHC and lipid antigens via CD1d pathways^16^. Moreover, γδ T cells can act as professional antigen-presenting cells in priming αβ T cells^17^ and support humoral immunity by promoting follicular helper T cell differentiation^18^. By expressing multiple chemokine receptors, γδ T cells home efficiently to inflamed tissues. Functionally, they release proinflammatory cytokines and cytotoxic mediators that accelerate inflammation and cell death and secrete chemokines that promote further recruitment of inflammatory and cytotoxic cells. Despite these pivotal roles, their contribution to atherogenesis and plaque destabilisation remains underexplored.

γδ T cells have been identified in chronically inflamed non-lymphoid tissues, such as the diabetic pancreas^19^, developing tumors^20^, and inflamed joints^21^, where they establish long-lived, tissue-resident memory populations^22^ that maintain homeostasis and immunosurveillance, functions likely relevant to chronic arterial inflammation. Although γδ T cells were first detected in human atherosclerotic plaques in 1993^23^, animal studies probing their role in atherosclerosis have produced conflicting results: one report implicated them in lesion development^24^, whereas another found them dispensable for plaque formation^25^. Further investigation is required to elucidate their precise contributions to atherogenesis and to inform potential therapeutic strategies.

In our study, we demonstrate that bone marrow (BM)-derived CD27+ γδ T cells exert potent atherogenic effects in experimental atherosclerosis. We identified two γδ T cell–specific effector molecules, interferon-γ (IFN-γ) and perforin (Pfp), as critical mediators of unstable plaque formation. As early responders to inflammation, these cells play a pivotal role in recruiting and modulating other immune cells, thereby amplifying localized inflammatory responses and promoting lesion necrosis. Exploiting the unique upregulated expression of CXCR3 on hyperlipidemic γδ T cells, we treated mice with established atherosclerosis using a small-molecule CXCR3 antagonist, AMG487. It significantly slowed lesion progression and enhanced plaque stability by reducing the infiltration of γδ T cells and other immune subsets and by diminishing lesion cell death. These results suggest that combining CXCR3 blockade strategy with conventional lipid-lowering therapies, such as statins, may synergistically prevent the development of vulnerable plaques and thereby reduce the risk of MI and stroke.

## Method

### Animals, ethics, diets and antibodies

All mice were maintained on a C57BL/6 background. Ldlr⁻/⁻ and TCRδ⁻/⁻ mice were obtained from Jackson Laboratory (Bar Harbor, ME, USA). Ifng⁻/⁻ and Prf1⁻/⁻ mice were kindly provided by Mark Smyth (QIMR Berghofer, Brisbane, Australia), and wild-type (WT) controls were sourced from the Walter and Eliza Hall Institute (Melbourne, Australia). All animal protocols were approved by the Alfred Research Alliance Animal Ethics Committee and carried out at the Precinct Animal Centre, Baker Heart and Diabetes Institute (Melbourne, Australia).

To induce lipid-driven atherosclerosis, mice were fed a HFD containing 0.15% cholesterol and 21% fat (Specialty Feeds, Glen Forrest, WA) with ad libitum access to sterile water and maintained on a 12-hour light/dark cycle. Antibodies used in the study were obtained from BD Biosciences based in San Jose, CA, and chemicals were sourced from Sigma-Aldrich in St. Louis, MO, unless otherwise specified.

### Generation of chimeric mice

To generate Ldlr⁻/⁻ mice devoid of γδ T cells (γδT^Deficient^), we employed total-body irradiation and BM transplantation as previously described^26^. Briefly, 6-week-old male Ldlr⁻/⁻ mice received divided total-body irradiation of 1,100Gy and were reconstituted with 5×10^6^ BM cells from TCRδ⁻/⁻ donors. Control Ldlr⁻/⁻ mice (γδT^Present^) received 5×10^6^ BM cells from WT donors. To investigate the contribution of γδ T cell–derived IFN-γ, we generated mixed BM chimeras^26,27^. Irradiated Ldlr⁻/⁻ recipients were transplanted with 4×10^6^ TCRδ⁻/⁻ plus 1×10^6^ IFN-γ⁻/⁻ BM cells, yielding mice lacking IFN-γ specifically in γδ T cells (IFN-γ-deficient γδT). Control chimeras (IFN-γ-competent γδT) received 4×10^6^ WT plus 1×10^6^ IFN-γ⁻/⁻ BM cells. An identical approach was used to generate Pfp-deficient γδ T cell chimeras, using Pfp^-/-^ donor mice.

### γδ T cell repletion and depletion

Spleen γδ T cells were isolated using the Miltenyi Biotec magnetic γδ T Cell Isolation Kit according to the manufacturer’s instructions. Cells were then stained with Streptavidin-APC and anti-mouse CD27-PE and sorted into CD27+ and CD27⁻ subsets (purity >99%) on a BD FACS Aria cell sorter. Viability (>95%) was confirmed before further use. For depletion studies, Ldlr⁻/⁻ mice with established atherosclerosis received 10 mg/kg of either anti-mouse TCR γ/δ antibody (UC7-13D5, BioXCell) or non-immune IgG (BE0091, BioXCell) via tail-vein injection every two week.

### Induction of tandem stenosis (TS) in the right common carotid artery

Atherosclerotic Ldlr⁻/⁻ mice were anesthetized by intraperitoneal injection of ketamine (80 mg/kg), xylazine (20 mg/kg), and atropine (0.06 mg/kg). Under sterile conditions, the right common carotid artery was exposed through a midline neck incision. Two sequential stenoses were created at 1 mm and 4 mm proximal to the carotid bifurcation using a custom constriction device (150 µm outer diameter), following our previously described protocol^28^. The device was carefully slid around the artery to produce consistent luminal narrowing. The incision was then closed in layers, and animals were allowed to recover under a heat lamp with postoperative monitoring until ambulatory.

### Statistical analysis

Statistical analyses were conducted using GraphPad Prism 7.04 software. Raw data were subjected to statistical tests including Student’s t-tests, Mann-Whitney U tests, or One-way analysis of variance (ANOVA) followed by Dunnett’s multiple comparison test or Kruskal-Wallis tests with Dunns post-test, depending on the distribution of the data. The results are presented as mean ± standard error of mean (SEM), and a p-value of less than 0.05 was considered statistically significant.

## Results

### γδ T cells contribute to atherosclerosis development

The presence of γδ T cells in human and murine atherosclerotic plaques is well documented in previous studies, which describe their accumulation in inflamed lesions and association with advanced disease stages^23–25^. In agreement with these reports, we confirmed the presence of γδ T cells in atherosclerotic lesions of the aortic sinus, innominate arteries, and aortic arch in Ldlr⁻/⁻ mice fed a HFD for 20 weeks (**Figure 1A**). To determine the functional role of γδ T cells in atherosclerosis, lethally irradiated Ldlr⁻/⁻ mice were reconstituted with TCRδ⁻/⁻ bone marrow to achieve γδ T cell depletion. Flow cytometry and immunofluorescence confirmed the effective absence of circulating and lesional γδ T cells in γδT^Deficient^ mice (**Figure 1B-C**), which was associated with an approximately 65% reduction in aortic plaque area compared to γδT^Present^ controls (**Figure 1D**). This reduction occurred despite comparable body weight and plasma cholesterol levels between the groups (**Figure 1E-F**), indicating a γδ T cell-specific effect on atherosclerosis. The diminished lesion size in γδT^Deficient^ mice was accompanied by significantly reduced infiltration of inflammatory and cytotoxic immune cells, including macrophages, CD4 T cells, CD8 T cells, and NK/NKT cells (**Figure 1G-H**). Moreover, these mice exhibited significantly smaller necrotic cores (**Figure 1I**) and reduced apoptotic cell burden (**Figure 1J**), indicative of a more stable plaque phenotype. Collectively, these findings highlight a central role for BM -derived γδ T cells in modulating intraplaque immune cell composition and necrosis, critical factors contributing to plaque progression and instability.

**Figure 1.**
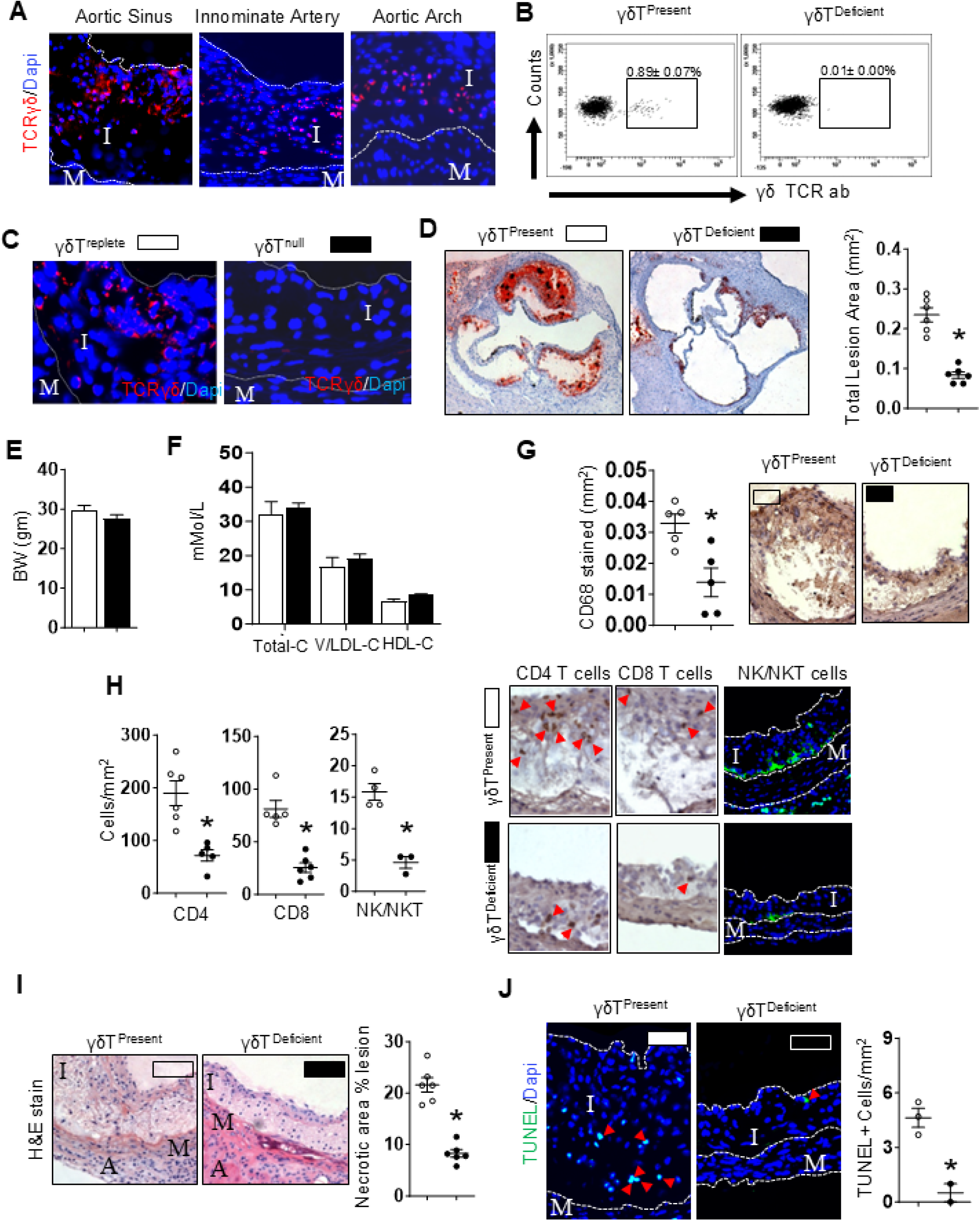
Chimeric depletion of γδ T cells reduces atherosclerotic burden and is associated with fewer inflammatory infiltrates, less apoptosis, and reduced necrosis. Atherosclerotic lesions from 20-week high-fat diet–fed mice were stained for γδ T cells by immunofluorescence and counterstained with DAPI. Fluorescent imaging revealed γδ T cells within the aortic sinus, innominate artery, and aortic arch lesions (**A**). γδT^Deficient^ Ldlr-/-mice lacked γδ T cells in the spleen (**B**) and within atherosclerotic plaques (**C**) and exhibited reduced lesion area (**D**). γδ T cell deficiency did not affect body weight (**E**) or plasma lipid cholesterol levels (**F**). Reduced lesion burden was accompanied by decreased infiltration of CD68-positive macrophages (**G**), CD4-positive, CD8-positive, and NK1.1-positive lymphocytes (**H**), as well as necrosis (**I**) and reduced apoptosis (**J**). Data are presented as mean ± SEM from two to three independent experiments and small circles representing data from individual mice. Representative micrographs and flow cytometry plots are shown. *p < 0.05. □: Ldlr-/-mice competent in γδ T cells (γδT^Present^); ▪: Ldlr-/-mice deficient in γδ T cells (γδT^Deficient^).

### CD27-positive γδ T cells promote atherosclerosis

To define the functional relevance of γδ T cell subsets in atherogenesis, we investigated CD27+ and CD27-γδ T cells, given that CD27 distinguishes two phenotypically and functionally distinct γδ T cell populations^29^. FACS-purified CD27-positive or CD27-negative γδ T cells (**Figure 2A**) were adoptively transferred into γδT^Deficient^ mice at weeks 0 and 4 of HFD feeding (1x10^5^ cells via tail vein injection). Flow cytometry at the end of 8 weeks confirmed effective reconstitution of γδ T cells in recipient mice (**Figure 2B**), with no significant differences in body weight or plasma cholesterol between groups (**Figure 2C-D**). Mice receiving CD27-positive γδ T cells developed significantly larger aortic plaques compared to those receiving CD27-negative γδ T cells or vehicle (**Figure 2E**). CD27 expression functionally segregates γδ T cells into CD27-positive IFN-γ–producing and CD27-negative IL-17– producing subsets^29^. Immunofluorescence analysis revealed markedly increased IFN-γ protein levels in atherosclerotic plaques of CD27-positive γδ T cell recipients compared to those receiving CD27-negative cells (**Figure 2F**). This was supported by gene expression analysis showing reduced plaque IFN-γ mRNA in γδT^Deficient^ mice (**Figure 2G**), which was restored upon reconstitution with CD27-negative γδ T cells (**Figure 2H**). In contrast, IL-17 protein and mRNA levels remained unchanged regardless of γδ T cell depletion or repletion (**Figure 2F–H**). Together, these findings indicate that CD27-positgive γδ T cells contribute to atherogenesis via IFN-γ– mediated manner, whereas IL-17–producing CD27-negative γδ T cells appear to be dispensable in this setting.

**Figure 2.**
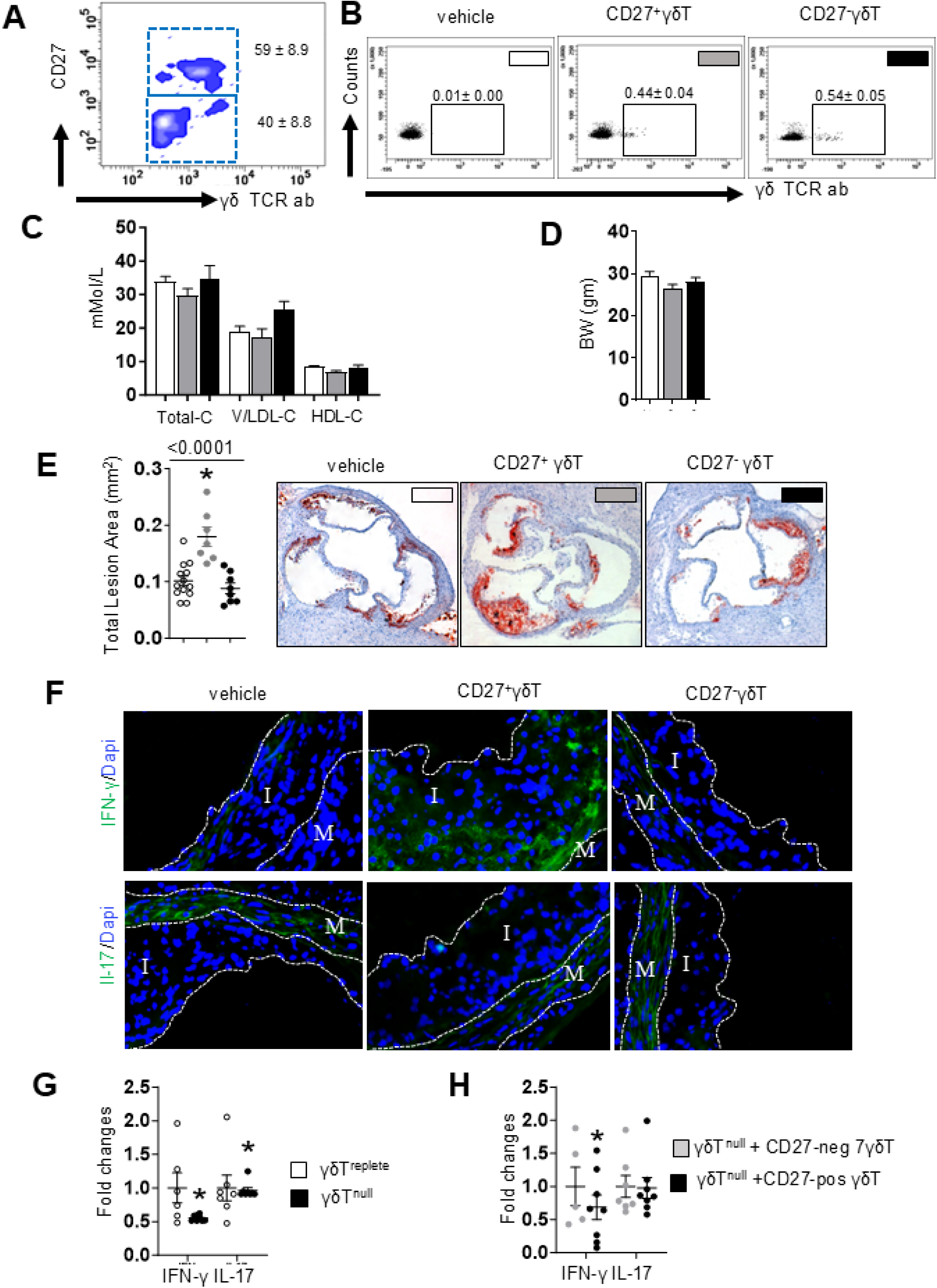
CD27-positive, but not CD27-negative, γδ T cells promote atherosclerosis. γδT^Deficient^ mice were adoptively transferred with either CD27-positive or CD27-negative γδ T cells isolated from wild-type spleens (**A**) at the start of an 8-week high-fat diet. At endpoint, transferred γδ T cells were detected in recipient mice, compared to vehicle controls (**B**), and no significant differences in plasma lipid profiles (**C**) or body weights (**D**) were observed among groups. However, mice that received CD27-positive γδ T cells exhibited significantly increased atherosclerosis compared to those receiving CD27-negative γδ T cells or vehicle alone (**E**). Immunofluorescent staining revealed upregulated IFN-γ protein expression in lesions following CD27-positive γδ T cell transfer, while IL-17 expression remained unchanged regardless of the γδ T cell subset transferred (**F**). These protein findings were supported by mRNA analysis, showing increased IFN-γ transcript levels only in mice receiving CD27-positive γδ T cells, whereas IL-17 mRNA levels were unaffected (**G–H**). Data are presented as mean ± SEM from two to three independent experiments and small circles representing data from individual mice. Representative immunofluorescence images and FACS plots are shown. *p < 0.05. □: γδT^0^ mice with vehicle; ▪: γδT^0^ mice with CD27+ γδ T cells; ▪: γδT^0^ mice with CD27-γδ T cells.

### γδ T cells are activated in atherosclerotic plaques

Cytotoxic responses and IFN-γ production by activated γδ T cells are mediated by killer-activating receptors such as CD27 and DNAM-1^15,30^, and are further enhanced by NKG2D- and CD16-dependent mechanisms^15,30^. Given the abundant presence of γδ T cells in both murine and human atherosclerotic plaques (**Figures 1A and 3A**), we next investigated whether lesional γδ T cells are functionally activated within plaques to exert pathogenic effector functions. Consistent with previous reports, immunofluorescence staining of sinus lesions from 16-week HFD-fed mice revealed the presence of CD70, CD155, RAE-1, and IgG within the plaques. Notably, lesional γδ T cells expressed the corresponding receptors, CD27, DNAM-1, NKG2D, and CD16, suggesting their potential engagement and activation within the plaque microenvironment (**Figure 3B**). These findings prompted us to assess the temporal dynamics of γδ T cell–activating ligand expression during plaque progression. Consistent with increasing plaque complexity, expression and accumulation of these activating ligands progressively increased over time. Notably, IgG emerged as the most abundantly deposited ligand in the atherosclerotic plaques (**Figures 3C and S1**). In normal adult mice, Vγ1⁺ γδ T cells represent a major population within peripheral lymphoid organs^31^, and CD27⁺ γδ T cells are predominantly composed of Vγ1⁺ subsets^32^. Based on this, we examined the distribution of Vγ1⁺ γδ T cells in atherosclerotic mice and found that Vγ1⁺ and Vγ2⁺ subsets together accounted for approximately 76% of CD27⁺ γδ T cells (**Figure 3D**). Given that both macrophages and CD27-positive γδ T cells, key innate immune responders, infiltrate atherosclerotic plaques early, prior to the accumulation of adaptive immune products such as IgG, we hypothesized that macrophages may promote CD27-positgive γδ T cell activation through co-stimulatory signals such as CD70 and CD155, as activated macrophages express these proteins under stress or inflammatory conditions Immunofluorescence analysis of aortic lesions from 2-, 4- and 6-week HFD-fed mice showed that lesional macrophages express CD70 and CD155, ligands for CD27 and DNAM-1 respectively (**Figures 3E-F and S2**). Together, these findings suggest that macrophages engage γδ T cells through the CD27–CD70 and DNAM-1–CD155 axes, promoting their effector activation during the early stages of plaque development, while RAE-1 and IgG become prominent at advanced stages.

**Figure 3.**
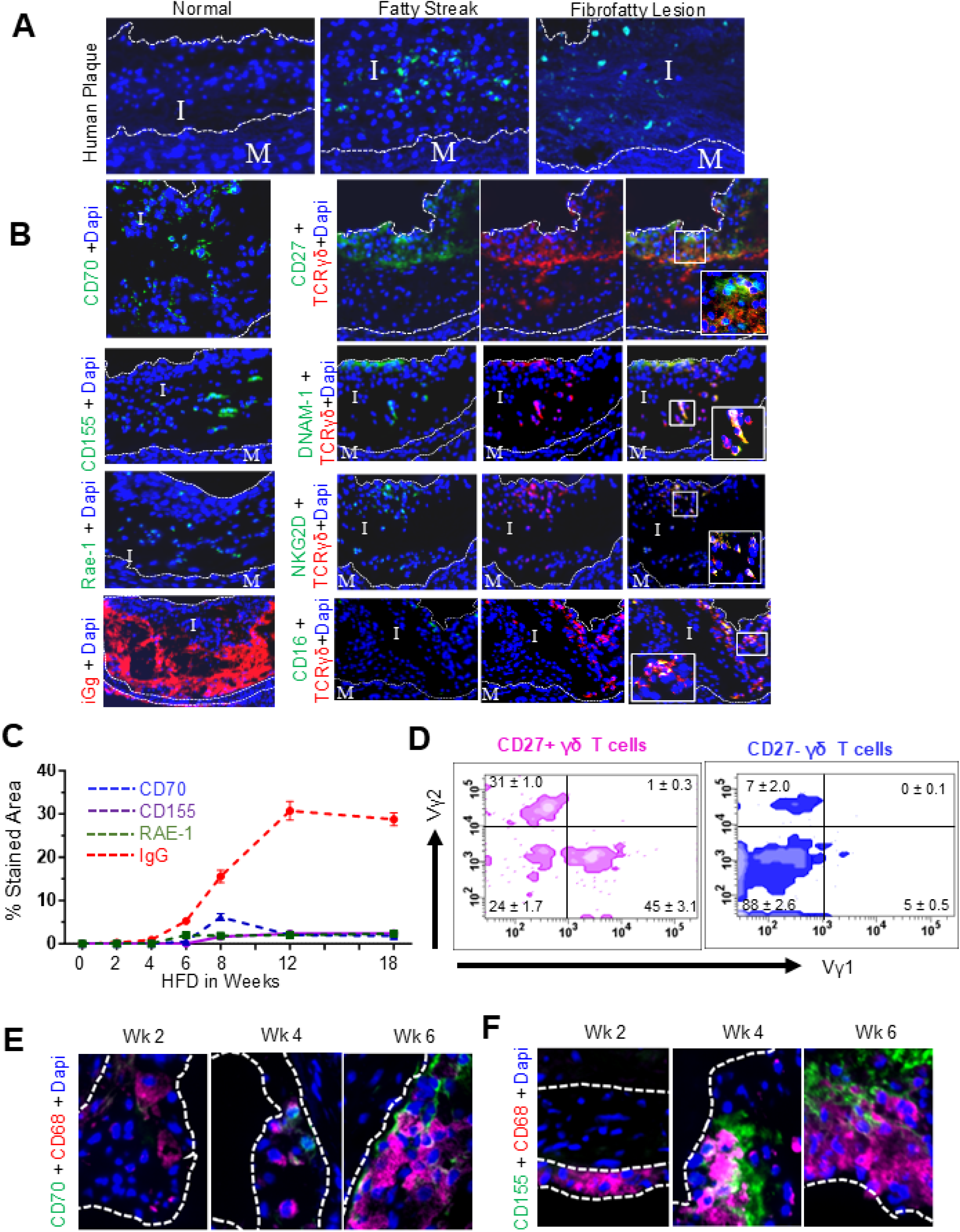
Protein ligands that activate γδ T cells are expressed in atherosclerotic plaques. Human atherosclerotic plaques were assessed for the presence of γδ T cells. Both fatty streaks and fibrofatty lesions demonstrated γδ T cell infiltration compared to normal tunica intima (**A**). Murine atherosclerotic lesions were immunofluorescently stained for protein ligands known to activate γδ T cells upon recruitment to plaques. Fluorescent imaging revealed the presence of CD70, CD155, RAE-1, and IgG, ligands for the γδ T cell-expressing receptors CD27, DNAM-1, NKG2D, and CD16, respectively (**B**). Temporal expression of these ligands within plaques was evaluated, showing that IgG deposition occurred later but was the most abundant among the detected ligands (**C**). CD27⁺ and CD27⁻ γδ T cell subsets were characterized using Vγ1 and Vγ2 antibodies (**D**). Macrophage expression of CD70 and CD155 was examined at different time points during lesion progression (**E-F**). Representative immunofluorescence images and FACS dot plot are shown; n = 4–8 mice per group.

### γδ T cell–derived IFN-γ initiates and propagates lesion inflammation

γδ T cells are potent producers of IFN-γ, particularly during early immune responses to pathogens and tumors^33^. To determine whether this effector function contributes to atherosclerosis, we first assessed IFN-γ expression by plaque-resident γδ T cells. Immunofluorescence colocalization studies revealed that approximately 85% of lesional γδ T cells in mice fed a HFD for 23 weeks expressed IFN-γ (**Figure 4A**), consistent with the functional dominance of the CD27-positive subset^29^. These findings suggest that IFN-γ–producing γδ T cells, rather than IL-17–producing subsets, are key drivers of plaque progression in this model. To directly assess the pathogenic role of γδ T cell–derived IFN-γ in atherosclerosis, we generated chimeric mice selectively lacking IFN-γ in γδ T cells. Flow cytometric analysis confirmed comparable γδ T cell numbers between test and control mice (**Figure 4B**), with selective loss of IFN-γ production in the γδ T cell compartment of test mice (**Figure 4C**). Despite similar plasma lipid levels and body weight (**Figure 4D–E**), mice lacking γδ T cell–derived IFN-γ developed significantly smaller atherosclerotic plaques compared to IFN-γ–competent controls (**Figure 4F**). Consistent with a central role for IFN-γ in orchestrating vascular inflammation, aortic mRNA expression of pro-inflammatory cytokines, including IFN-γ, TNF-α, IL-1β, IL-18, IL-12, and IL-15, was markedly reduced in IFN-γ–deficient γδT mice (**Figure 4G**). In addition, these mice exhibited smaller necrotic cores and reduced lesional apoptosis (**Figure 4H–I**), implicating γδ T cell–derived IFN-γ as a key mediator of plaque progression, vascular inflammation, and inflammation-induced tissue damage.

**Figure 4.**
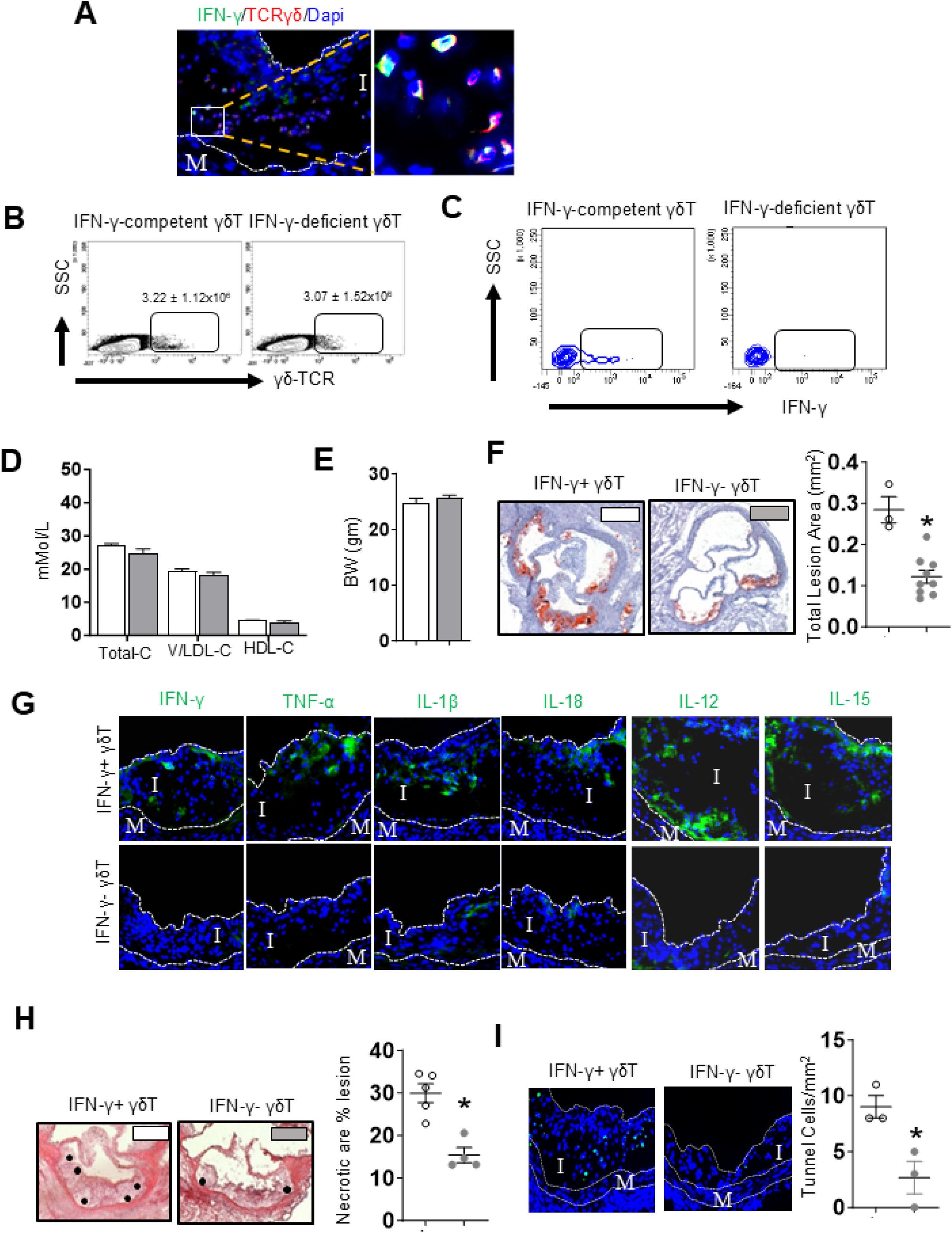
IFN-γ derived from γδ T cells is critically important for plaque development. Immunofluorescent staining identified IFN-γ-producing γδ T cells within atherosclerotic lesions of 23-week high-fat diet-fed Ldlr⁻/⁻ mice (**A**). Chimeric Ldlr⁻/⁻ mice selectively deficient in IFN-γ in γδ T cells were generated and confirmed via flow cytometry. FACS analysis demonstrated comparable γδ T cell numbers between IFN-γ-deficient and control mice (**B**), but γδ T cells from IFN-γ-deficient mice failed to produce IFN-γ upon stimulation, in contrast to αβ T cells (**C**). Plasma cholesterol levels and body weights were comparable across groups (**D-E**). Mice lacking IFN-γ in γδ T cells exhibited significantly reduced atherosclerotic burden (**F**). This reduction was accompanied by lower lesional expression of proinflammatory cytokines including IFN-γ, TNF-α, IL-1β, IL-18, IL-12, and IL-15 (**G**) Lesions from IFN-γ-deficient γδ T cell mice showed reduced necrotic core areas (**H**) and decreased numbers of TUNEL-positive cells (**I**), indicating reduced cell death. Data are presented as mean ± SEM from two to three independent experiments and small circles representing data from individual mice. Representative immunofluorescence images and FACS dot plots are shown. p < 0.05. □: IFN-γ-competent γδT; ▪: IFN-γ-deficient γδT T cells.

### γδ T cells induce direct lesion cell death via a perforin-dependent mechanism

γδ T cells can eliminate target cells via multiple cytotoxic mechanisms, including Pfp-mediated granule exocytosis, Fas/FasL–induced caspase-8–dependent apoptosis, and TRAIL-induced extrinsic apoptosis. To determine whether lesional γδ T cells express these death-inducing molecules, we performed immunofluorescence staining. This revealed high levels of perforin expression (>80%) in lesional γδ T cells, whereas expression of FasL and TRAIL was minimal (<8–10%) (**Figure 5A**), suggesting that granule-mediated killing is the dominant effector pathway, consistent with previous findings^34^. CD27-positive γδ T cells, a subset known for producing IFN-γ, have also been reported to express high levels of perforin^35^. To assess whether this cytotoxic subset contributes to atherosclerosis, we generated mixed bone marrow chimeras in which only γδ T cells lacked Pfp. These mice exhibited comparable γδ T cell numbers (**Figure 5B**) but lacked γδ T cell–specific perforin expression (**Figure 5C**). Despite having similar body weight (**Figure 5D**) and plasma cholesterol levels (**Figure 5E**) to controls, they developed significantly smaller atherosclerotic plaques (**Figure 5F**), accompanied by reduced necrotic core areas (**Figure 5G**) and fewer TUNEL+ apoptotic cells (**Figure 5H**). These findings indicate that γδ T cell–mediated cytotoxicity via perforin directly contributes to cell death and plaque destabilization. To rule out compensatory upregulation of alternative death pathways, we examined the expression of Fas, TRAIL, and FcγRIII (CD16) in the plaques and found no significant differences between groups (**Figure 5I**). Despite unchanged expression of these pathways, the substantial reduction in necrotic core size in mice lacking γδ T cell-derived perforin underscores γδ T cell–derived cytotoxins as pivotal role in γδ T cell–mediated atherogenesis.

**Figure 5.**
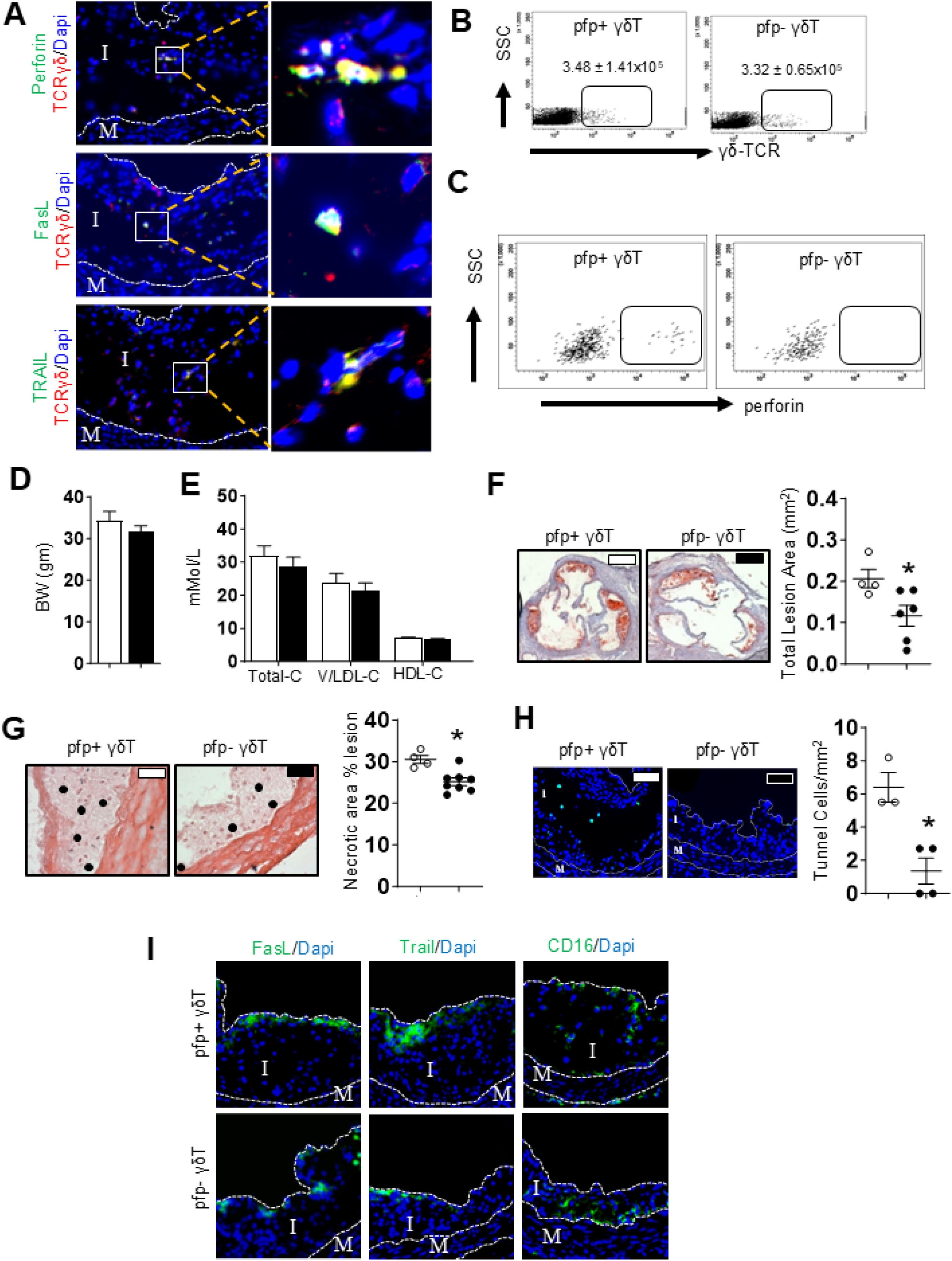
Cytotoxin-mediated cytotoxicity is essential for the effector function of γδ T cells in promoting lesion development. Immunofluorescent staining of atherosclerotic lesions from 23-week high-fat diet-fed Ldlr⁻/⁻ mice revealed that lesional γδ T cells express perforin (Pfp), Fas ligand (FasL), and TRAIL (**A**). Splenocytes from chimeric Ldlr⁻/⁻ mice selectively deficient in Pfp within γδ T cells were analysed using FACS. Although γδ T cell numbers remained unchanged (**B**), Pfp-deficient γδ T cells failed to express perforin upon stimulation, in contrast to NK1.1⁺ NK/NKT cells (**C**). Pfp deficiency in γδ T cells did not alter body weight (**D**) or plasma cholesterol levels (**E**). However, mice with Pfp-deficient γδ T cells exhibited significantly reduced atherosclerotic lesion size (**F**). These mice also showed reduced necrosis and fewer TUNEL⁺ apoptotic cells within lesions (**G-H**). Despite Pfp deficiency, lesional γδ T cells still expressed FasL, TRAIL, and CD16, suggesting additional cytotoxic mechanisms may remain active (**I**). Data are presented as mean ± SEM from two to three independent experiments and small circles representing data from individual mice. Representative immunofluorescence images nd FACS plots are shown. p < 0.05. □: Pfp -competent γδT; ▪: Pfp -deficient γδT T cells

### γδ T cells act as master regulators of plaque inflammation and cellular composition

Circulating γδ T cells are among the earliest immune responders recruited to sites of tissue injury, infection, or inflammation. They secrete a broad array of chemoattractants that amplify the immune response by promoting the recruitment of additional γδ T cells and a diverse range of myeloid and lymphoid effector cells. In addition to serving as an early and potent source of IFN-γ^33^, neuroinflammatory γδ T cells produce high levels of RANTES and MIP-1α, two key chemokines involved in immune cell recruitment^36^. Targeting IFN-γ and these IFN-γ-inducible chemoattractants in atherosclerotic mice has been shown to significantly reduce the recruitment of both myeloid and lymphoid cells to atherosclerotic lesions^37–39^. In accordance with these findings, we hypothesized that γδ T cells orchestrate broader immune activation within atherosclerotic plaques. We observed robust expression of the pro-inflammatory cytokine IFN-γ (**Figure 4A**), as well as the chemokines RANTES and MIP-1α (**Figure 6A-B**), in lesional γδ T cells, suggesting an activation loop driven by local stress signals. To test whether γδ T cells are required to sustain and propagate plaque inflammation, we depleted γδ T cells in mice with established atherosclerosis. Antibody-mediated depletion of γδ T cells led to a marked reduction in lesional macrophages, T cells, and NK/NKT cells (**Figure 6C**). Importantly, lesion size, necrotic core area, and apoptotic cell burden were all significantly reduced (**Figure 6D-F**), indicating that γδ T cells are critical for maintaining the inflammatory and cytotoxic environment that drives lesion progression. Based on our findings and prior reports showing that IFN-γ receptor (IFNGR) deficiency reduces lesion size and alters plaque composition in atherosclerotic mice composition^40^, we hypothesized that γδ T cell–derived IFN-γ plays a central role in amplifying plaque progression through self-sustaining inflammatory loops. Supporting this, we found that cytotoxic and inflammatory lymphocyte populations within the plaque expressed IFNGR (**Figure 6G**), indicating their capacity to respond to γδ T cell–derived IFN-γ. Collectively, these data reveal a pivotal role for γδ T cell–derived IFN-γ in promoting atherosclerosis through both autocrine and paracrine mechanisms.

**Figure 6.**
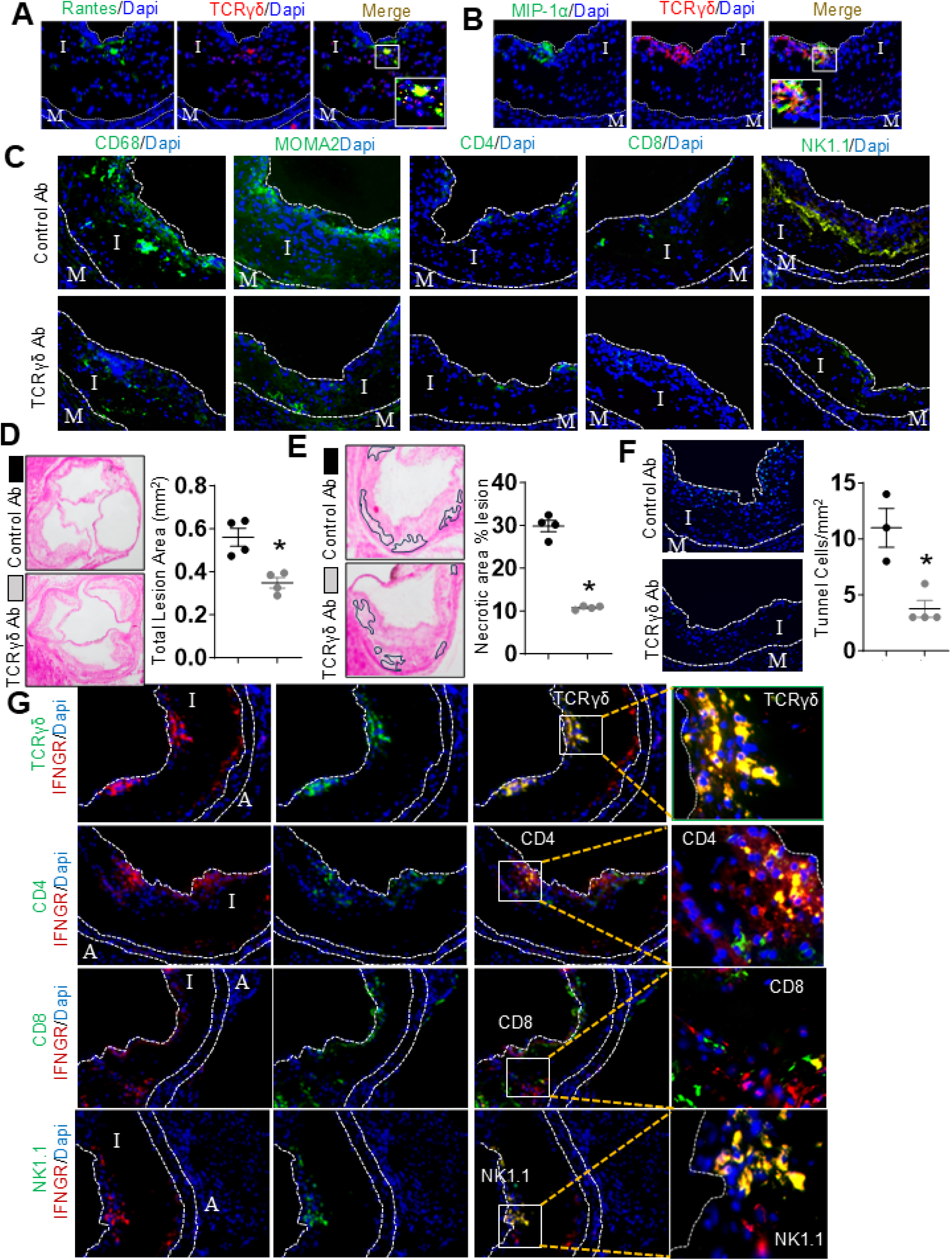
Actively recruited γδ T cells play a crucial role in plaque progression. Immunofluorescent staining of atherosclerotic lesions from 23-week high-fat diet-fed Ldlr⁻/⁻ mice demonstrated that lesional γδ T cells express the potent chemoattractants RANTES and MIP-1α, which are primarily involved in monocyte and lymphocyte recruitment (**A-B**). Ldlr⁻/⁻ mice with established atherosclerosis were treated with a γδ T cel-depleting antibody. Antibody treatment led to efficient depletion of lesion γδ T cells and was also associated with reduced accumulation of other immune cells in lesions, including CD68⁺ macrophages, MOMA⁺ monocytes/macrophages, and CD4⁺, CD8⁺, and NK1.1⁺ lymphocytes (**C**). Depletion of γδ T cells resulted in reduced atherosclerosis progression (**D**), accompanied by decreased lesional necrosis and apoptosis (**E-F**). Immunofluorescence imaging further confirmed the expression of IFN-γ receptor (IFNGR) on lesional TCRγδ⁺, CD4⁺, CD8⁺, and NK1.1⁺ immune cells in atherosclerotic lesions from 23-week HFD-fed *Ldlr⁻/⁻* mice (**G**). Data are presented as mean ± SEM from two to three independent experiments and small circles representing data from individual mice. Representative immunofluorescence images are shown. *p* < 0.05. □: control antibody treated mice; ▪: γδ T cell depleting antibody treated mice.

### γδ T Cells accumulate over time in atherosclerotic lesions

Given our findings that γδ T cells contribute to plaque instability and potentially plaque rupture, we next examined their presence in unstable atherosclerotic plaques from preclinical tandem-stenosis mice. Immunostaining of adjacent sections revealed abundant γδ T cells at sites of plaque rupture (**Figure 7A**). Consistent with their progressive accumulation during lesion development (**Figure 7B**) and increased expression of CD69 and CD103 in rupture-prone plaques (**Figure 7C**), these data suggest that γδ T cells infiltrate atherosclerotic lesions and may acquire a tissue-resident memory phenotype. This phenotype likely contributes to local immune activation and instability, providing a mechanistic explanation for their critical role in plaque destabilization. To prevent γδ T cell trafficking to atherosclerotic lesions, which is likely mediated by chemokine -chemokine receptor interactions, we first characterized their chemokine receptor expression profile under hyperlipidemic conditions as peripheral migratory γδ T cells dynamically alter their chemokine receptor repertoire in response to disease state, inflammation, and tissue context^41–44^.

**Figure 7.**
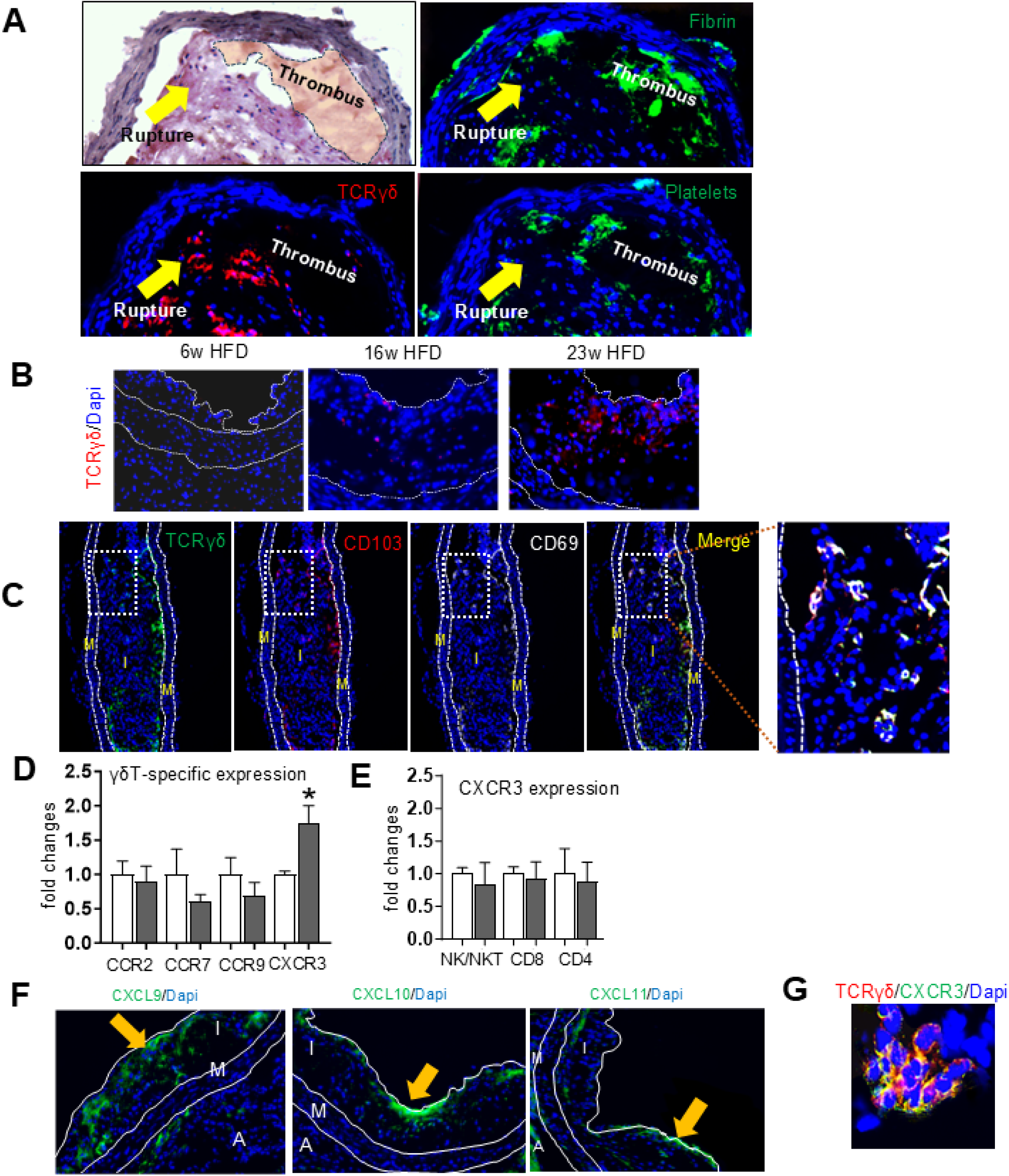
γδ T cells accumulate near plaque rupture sites. Carotid artery sections from tandem-stenosis atherosclerotic mice were examined using histology and immunofluorescence. Serial 6 μm sections revealed dense accumulation of γδ T cells at plaque rupture sites, characterized by disrupted endothelial layers and luminal thrombosis positive for fibrin and CD42 (**A**). Time-course fluorescent imaging of atherosclerotic lesions from Ldlr⁻/⁻ mice fed a high-fat diet for 6, 16, and 23 weeks showed progressive accumulation of γδ T cells within plaques (**B**). Co-immunofluorescence staining on carotid artery sections from tandem-stenosis atherosclerotic mice demonstrated colocalization of γδ T cells with CD69 and CD103, markers indicative of tissue-resident memory phenotype (**C**). γδ T cells were isolated from spleens of HFD-fed Ldlr⁻/⁻ mice and chow-fed WT mice. RT-PCR analysis showed a significant upregulation of CXCR3 in γδ T cells from hyperlipidemic mice (**D**), while expression levels of CXCR3 in NK1.1⁺, CD8⁺, and CD4⁺ cells remained unchanged (**E**). Protein expression of CXCR3 ligands (CXCL9, CXCL10, CXCL11) were upregulated in atherosclerotic aortas/plaques, with marked localization near the endothelial lining (**F**). CXCR3⁺ γδ T cells were also detected within atherosclerotic plaques (**G**). Representative microimages are shown; n = 4–5 mice per group. p < 0.05. □: hyperlipidemic mice; ▪: normolipidemic mice.

To assess this in a hypercholesterolaemic setting, we isolated splenic γδ T cells from hyperlipidaemic mice and performed RT-PCR analysis of chemokine receptor transcripts. CXCR3 emerged as the predominant receptor (**Figure 7D**), consistent with prior studies showing preferential homing of CXCR3⁺ γδ T cells to endothelial sites^43^. In contrast, other lymphocyte subsets resident in plaques, including CD4⁺, CD8⁺, and NK1.1⁺ cells, showed no upregulation of CXCR3 in hyperlipidaemic mice (**Figure 7E**). Consistent with reports documenting elevated expression of the IFN-γ– inducible chemokines CXCL9, CXCL10, and CXCL11 in both human and murine atherosclerotic plaques across disease stages^45,46^, we observed significantly increased protein expression of these ligands within the lesions (**Figure 7F**). Immunofluorescence further revealed that these chemokines were upregulated in areas adjacent to the endothelial layer (**Figure 7F**) and CXCR3⁺ γδ T cells within the plaques (**Figure 7G**). Moreover, γδ T cells were found in proximity to, and colocalized with, key pro-inflammatory mediators, including RANTES, MIP-1α, IFN-γ, and TNF-α, in coronary lesions (**Figure S3**). Collectively, these findings reveal a previously unrecognized role for γδ T cells in promoting plaque inflammation, instability, and rupture, highlighting their critical importance in coronary plaque rupture.

### CXCR3 antagonist prevents unstable plaque development

Our previous strategy to inhibit γδ T cell–mediated plaque progression involved both local and systemic depletion of these cells, which significantly promoted plaque stabilization (**Figure 4**). To achieve this effect pharmacologically, without broadly suppressing the immune system, we evaluated AMG 487, a small-molecule CXCR3 antagonist previously shown to inhibit CXCR3-dependent metastasis in cancer models^47,48^, for its potential to limit plaque progression and destabilization.

Hypercholesterolemic Ldlr⁻/⁻ mice with tandem stenosis were treated with AMG 487 for an additional eight weeks. AMG 487 treatment did not affect splenic γδ, CD4⁺, or CD8⁺ T cell numbers, body weight, or plasma cholesterol levels (**data not shown**). Despite this, AMG 487 significantly reduced atherosclerotic burden, as evidenced by decreased lesion size both macroscopically (**Figure 8A**) and microscopically (**Figure 8B**). As expected, treated mice exhibited not only attenuated plaque growth but also enhanced plaque stability, characterized by increased vascular smooth muscle content and cap thicker fibrous caps (**Figure 8C–E**). These effects were accompanied by reduced macrophage infiltration (**Figure 8F**) and decreased numbers of lesional γδ T cells, CD4⁺ and CD8⁺ T cells, and NK/NKT cells (**Figure 8G**). These cellular alterations were associated with smaller necrotic cores (**Figure 8H**), fewer TUNEL⁺ apoptotic cells (**Figure 8I**), and reduced intraplaque levels of pro-inflammatory cytokines including IFN-γ, TNF-α, IL-1β, IL-18, IL-15, and IL-12 (**Figure S4**), reflecting suppressed inflammation and cell death. Collectively, these findings suggest that pharmacological blockade of CXCR3 with AMG 487 effectively attenuates atherosclerosis progression and mitigates features of plaque vulnerability.

**Figure 8.**
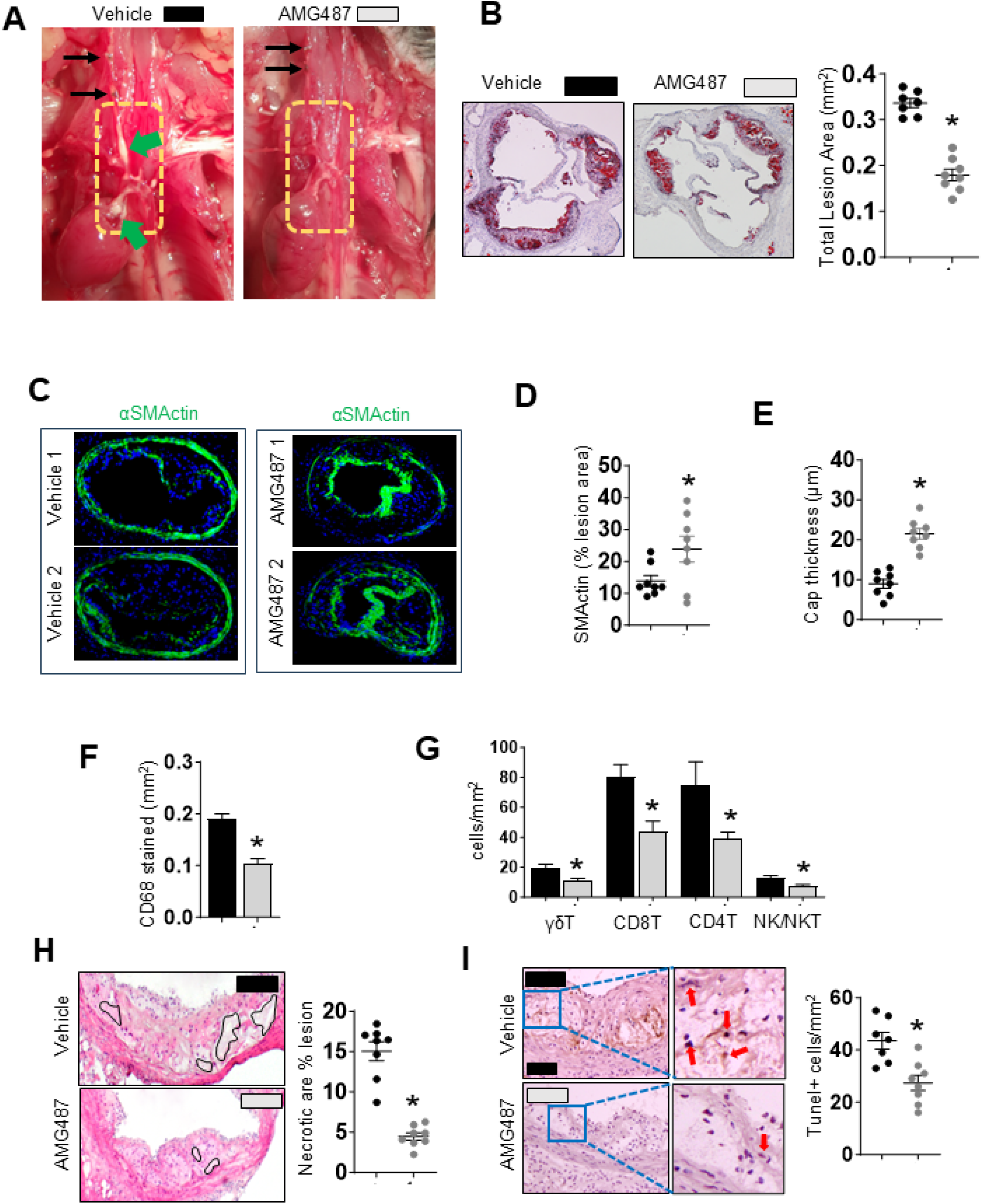
AMG487 treatment reduces plaque progression and enhances plaque stability. The therapeutic efficacy of the CXCR3 antagonist AMG487 was evaluated in hypercholesterolemic Ldlr⁻/⁻ mice subjected to carotid tandem-stenosis. AMG487 treatment significantly reduced atherosclerotic burden at the innominate artery (assessed macroscopically) and at the aortic sinus (**A-B**). Immunofluorescent staining of TS segment I revealed that treatment increased vascular smooth muscle content (α-SM actin-positive) in the intima and near endothelium (**C**). Treated mice showed significantly increased vascular smooth muscle content and fibrous cap thickness compared to vehicle controls (**D-E**). AMG487 also decreased lesional immune CD680positive macrophage and lymphocyte cell infiltration (**F-G**), necrotic core size (**H**) and apoptotic cell numbers (**I**). Data are presented as mean ± SEM from two to three independent experiments and small circles representing data from individual mice. Representative immunofluorescence images are shown. *p* < 0.05. ¢: Vehicle-treated mice; £: AMG487-treated mice. ▪: Vehicle treated mice; ▪: AMG487 treated mice.

## Discussion

In this study, we demonstrate, for the first time, that BM–derived, CD27-positive γδ T cells are critical drivers of atherosclerosis, directly and indirectly promoting plaque growth and instability. We also show that pharmacological blockade of CXCR3, a chemokine receptor highly enriched on hyperlipidemic γδ T cells, offers a promising strategy to limit γδ T cell-mediated plaque instability. This positions γδ T cells not merely as bystanders but as central orchestrators of vascular inflammation and lesion progression.

The developmental origin of γδ T cells dictates their effector cytokine profile, with most IL-17-producing γδ T cells arising from the embryonic thymus. In contrast, IFN-γ-producing γδ T cells are predominantly BM–derived and characteristically express CD27, a surface marker that distinguishes IFN-γ–producing (CD27-positive) from IL-17–producing (CD27-negative) subsets^29^. Consistent with previous reports showing that IL-17–expressing γδ T cells are virtually absent in chimeric mice reconstituted with BM-hemopoietic cells^49^, our findings highlight a key atherogenic role for BM-derived IFN-γ–producing, rather than thymus-derived IL-17–producing, γδ T cells.

This was demonstrated through loss-of-function experiments and further validated by adoptive transfer studies showing the pathogenic contribution of CD27-positive γδ T cells. We observed robust IFN-γ, but not IL-17 expression at both protein and mRNA levels within atherosclerotic lesions, with dynamic modulation in both depletion and reconstitution models. Furthermore, IFN-γ–deficient, not IL-17-deficient γδ T cells failed to promote plaque growth and instability, underscoring the critical role of IFN-γ derived from γδ T cells. Collectively, these data establish BM-derived CD27-positive γδ T cells as indispensable drivers of IFN-γ–mediated inflammation and atherosclerotic progression, leading to plaque instability.

Migratory γδ T cells function as rapid sentinels of infection and cellular stress, orchestrating all phases of local inflammation^50^. These cells are abundantly present in advanced human and murine atherosclerotic lesions and progressively accumulate in inflamed aortas in a time-dependent manner. Monocytes are the first immune cells to infiltrate the vascular wall, differentiating into macrophages, particularly pro-inflammatory M1 macrophages, in response to endothelial injury and the presence of oxidized LDL (oxLDL). These activated macrophages produce high levels of IFN-γ, which in turn induces expression of IFN-γ–responsive chemokines, CXCL9, CXCL10, and CXCL11, by both macrophages and endothelial cells^51^. It is plausible that hyperlipidaemia-induced CXCR3-positive γδ T cells infiltrate the arterial wall through interactions with the CXCR3–CXCL9/10/11 axis, a pathway active throughout all stages of atherosclerosis. This chemotactic axis, well-established in tumours and chronic inflammatory diseases, is now increasingly recognized as a central mediator of vascular immunopathology^51^. In line with previous studies reporting reduced plaque burden and inflammation in CXCR3-deficient mice^46^, our loss- and gain-of-function approaches, coupled with CXCR3 antagonism using AMG 487, provide an explanation governing their early recruitment as well as demonstrate the critical role of CD27-positive γδ T cells in both the initiation and progression of atherosclerotic plaque development.

IFN-γ is a well-established driver of plaque destabilization in atherosclerosis, acting through multiple downstream mechanisms. First, it induces the expression of chemokines such as CXCL9, CXCL10, and CXCL11, which recruit CXCR3-expressing monocytes, γδ T cells, and other immune cells to the lesion^51^, sustaining a self-amplifying inflammatory loop. Second, IFN-γ promotes the production of various proinflammatory cytokines, such as TNF-α, IL-1β, IL-12, IL-5, and IL-18^52^, thereby amplifying immune activation within the plaque microenvironment. Third, it induces multiple forms of programmed cell death, including direct apoptosis^53^ as well as synergistic forms such as PANoptosis, pyroptosis, and necroptosis, particularly when combined with co-stimulatory signals like TNF-α, pathogen-associated molecular patterns (PAMPs), and inflammasome activation^54^. Fourth, IFN-γ impairs collagen synthesis and vascular smooth muscle cell function^55^, weakening the fibrous cap and predisposing plaques to rupture. It suppresses the production of collagens I and III and promotes the expression of matrix metalloproteinases (MMPs) that degrade the extracellular matrix^39^. Fifth, IFN-γ enhances oxidative stress and endothelial dysfunction by upregulating inducible nitric oxide synthase (iNOS) and NADPH oxidase components, leading to the generation of reactive oxygen and nitrogen species^39^ that further damage vascular cells and oxidize lipids, thereby exacerbating plaque vulnerability. Literature demonstrates that deficiency of IFN-γ or IFNAR in atherosclerotic mice reduces atherosclerosis, accompanied by decreased inflammatory cell infiltration cells^40,56^. In line with these findings, exogenous administration of IFN-γ exacerbates atherosclerosis, leading to increased immune cell infiltration cells^57^, highlighting the critical role of IFN-γ not only in the initiation, but also in the progression and rupture of atherosclerotic plaques. Therapeutically, IFNGR-blocking antibodies targeting IFN-γ have been shown to attenuate disease progression^58^.

γδ T cells, as an abundant early source of IFN-γ^59^, can act on macrophages, other γδ T cells, conventional T cells, and NK/NKT cells via IFN-γ receptor (IFNGR) signaling to promote immune cell activation, proliferation, differentiation, and survival^60^. IFN-γ secretion by γδ T cells is potentiated by pro-inflammatory cytokines such as TNF-α, IL-12, IL-15, and IL-18, which are abundant in the plaque microenvironment^61^. Thus, these cytokines not only enhance IFN-γ output but also promote the recruitment and survival of other immune cells, thereby reinforcing the inflammatory milieu. Furthermore, IFN-γ enhances macrophage production of TNF-α through IFNGR signaling^62^, leading to synergistic inflammatory responses.

Collectively, γδ T cells not only produce large amounts of IFN-γ in the early phase of inflammation, but this also initiates cytokine crosstalk that drives lesional cell death, promotes necrotic core expansion, and increases the risk of plaque rupture Beyond their well-established capacity to secrete pro-inflammatory cytokines and chemokines, γδ T cells possess potent cytotoxic capabilities that critically contribute to plaque immunopathology. These cells are equipped for granule-mediated cytotoxicity through the release of perforin and granzymes, utilizing the γδ T cell receptor (TCR) in coordination with activating receptors, such as natural killer cell receptors (NKRs)^63^, to recognize and eliminate stressed or transformed vascular cells. In addition, activated γδ T cells express death-inducing ligands including Fas ligand (FasL) and TNF-related apoptosis-inducing ligand (TRAIL), which engage cognate receptors on target cells to induce apoptosis^63^. A subset of γδ T cells also expresses CD16 (FcγRIII), enabling antibody-dependent cellular cytotoxicity (ADCC) against opsonized targets^63^. Our data demonstrate that lesional γδ T cells actively engage cytotoxin-mediated mechanisms, underscoring their direct role in promoting cell death and plaque destabilization. Importantly, γδ T cells also propagate lesion inflammation and cytotoxicity indirectly by activating NK cells and CD8⁺ T cells, as well as by presenting antigens to αβ T cells to potentiate adaptive immune responses. Although numerically less abundant than macrophages or conventional αβ T cells within plaques, γδ T cells exert a disproportionately high pathogenic impact by bridging innate and adaptive immunity, amplifying local cytotoxic cascades, and thereby exacerbating plaque instability. These findings position γδ T cells as pivotal effectors in the immunopathogenesis of atherosclerosis and highlight their therapeutic potential as a target to stabilize vulnerable plaques.

Targeting CXCR3 interrupts the recruitment of CXCR3-expressing myeloid and lymphoid cells, including γδ T cells, to sites of vascular inflammation^64^, thereby breaking a pathogenic positive feedback loop that sustains immune activation. In both γδ T cell–depleted atherosclerotic mice and genetic knockout models, we identify CD27-positive γδ T cells as pivotal amplifiers of this loop, driving leukocyte influx, escalating cytokine production, and promoting tissue injury. Pharmacological inhibition of CXCR3 with AMG 487 markedly reduced lesional immune cell infiltration, IFN-γ and other proatherogenic cytokine expression, and multiple indices of plaque vulnerability, effects that occurred independently of lipid modulation, underscoring the immune-driven nature of these benefits. These findings highlight the therapeutic promise of targeting the CXCR3 axis to mitigate γδ T cell–mediated vascular inflammation and improve plaque stability.

Our study has several limitations that should be acknowledged. First, due to technical constraints, we analyzed the γδ T cell compartment as a whole, without resolving functionally distinct subsets. Future studies employing Vγ-chain–specific tools or lineage-tracing approaches will be essential to define the contributions of individual γδ T cell lineages. Second, we were unable to distinguish between newly recruited and locally expanded γδ T cells within atherosclerotic plaques. The dynamics of γδ T cell tissue residency, proliferation, and recirculation in vascular lesions remain poorly understood and warrant further investigation. Despite these limitations, our findings position γδ T cells as uniquely capable of sustaining inflammation, executing cytotoxic functions, and amplifying cytokine-driven cell death within plaques. Their expression of CXCR3, responsiveness to plaque-derived inflammatory signals, and potent effector repertoire underscore their role as an underrecognized but critical immune population in the pathogenesis of atherosclerosis.

In conclusion, BM-derived, CD27+ γδ T cells emerge as pivotal drivers of atherosclerosis. By homing to plaques via CXCR3, secreting IFN-γ, deploying perforin-mediated cytotoxicity, and amplifying chemokine-centred immune recruitment, they create a self-reinforcing inflammatory loop that enlarges lesions and promotes necrotic-core formation. Disabling γδ T-cell IFN-γ or perforin, depleting γδ T cells, or blocking their CXCR3-directed trafficking with AMG 487 all markedly reduced plaque burden and instability—even in the face of persistent hyperlipidaemia. Collectively, these findings position γδ T-cell trafficking and effector functions as attractive therapeutic targets for stabilising vulnerable atherosclerotic plaques.

## Data availability

The data underlying this article will be shared at reasonable request to the corresponding author.

---

**Supplementary Figure S1.**
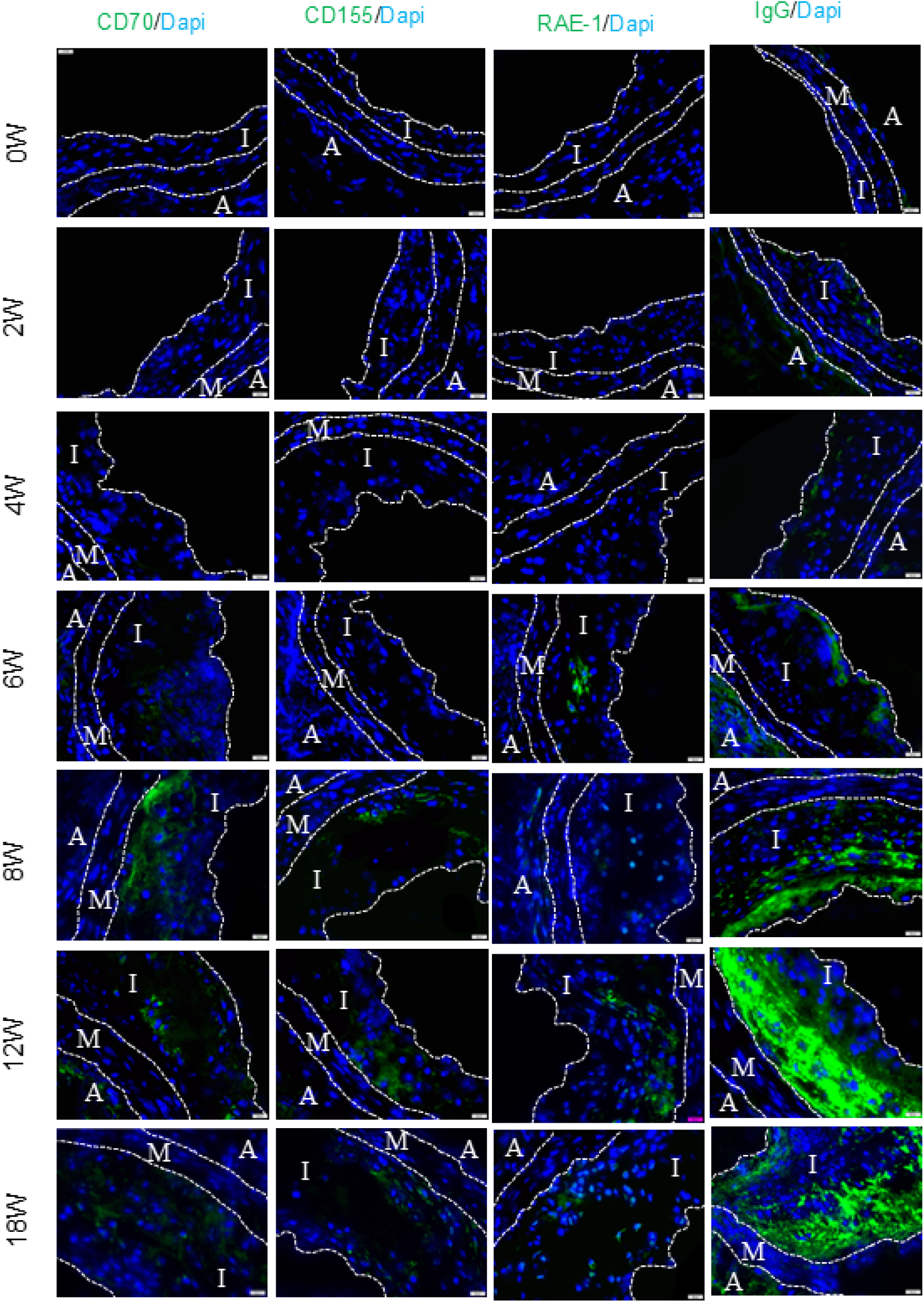
Longitudinal assessment of innate and adaptive death ligands in developing plaques. CD70, CD155, RAE-1, and IgG are stress-inducible ligands associated with innate immune responses and play key roles in mediating target cell death. In addition, Fc fragments of adaptive immune-derived IgGs serve as ligands that engage both innate and adaptive immune effector mechanisms, contributing to target cell death via antibody-dependent cell-mediated cytotoxicity (ADCC) and antibody-dependent cellular phagocytosis (ADCP). Fluorescent immunostaining was performed on atherosclerotic plaques from the aortic sinus collected at various time points. Imaging analysis revealed progressive upregulation of these death ligands during plaque development. Representative immunofluorescence images are shown; *n* = 5–8 mice per group. I: Tunica intima; M: Tunica media; A: Tunica adventitia.

**Supplementary Figure S2.**
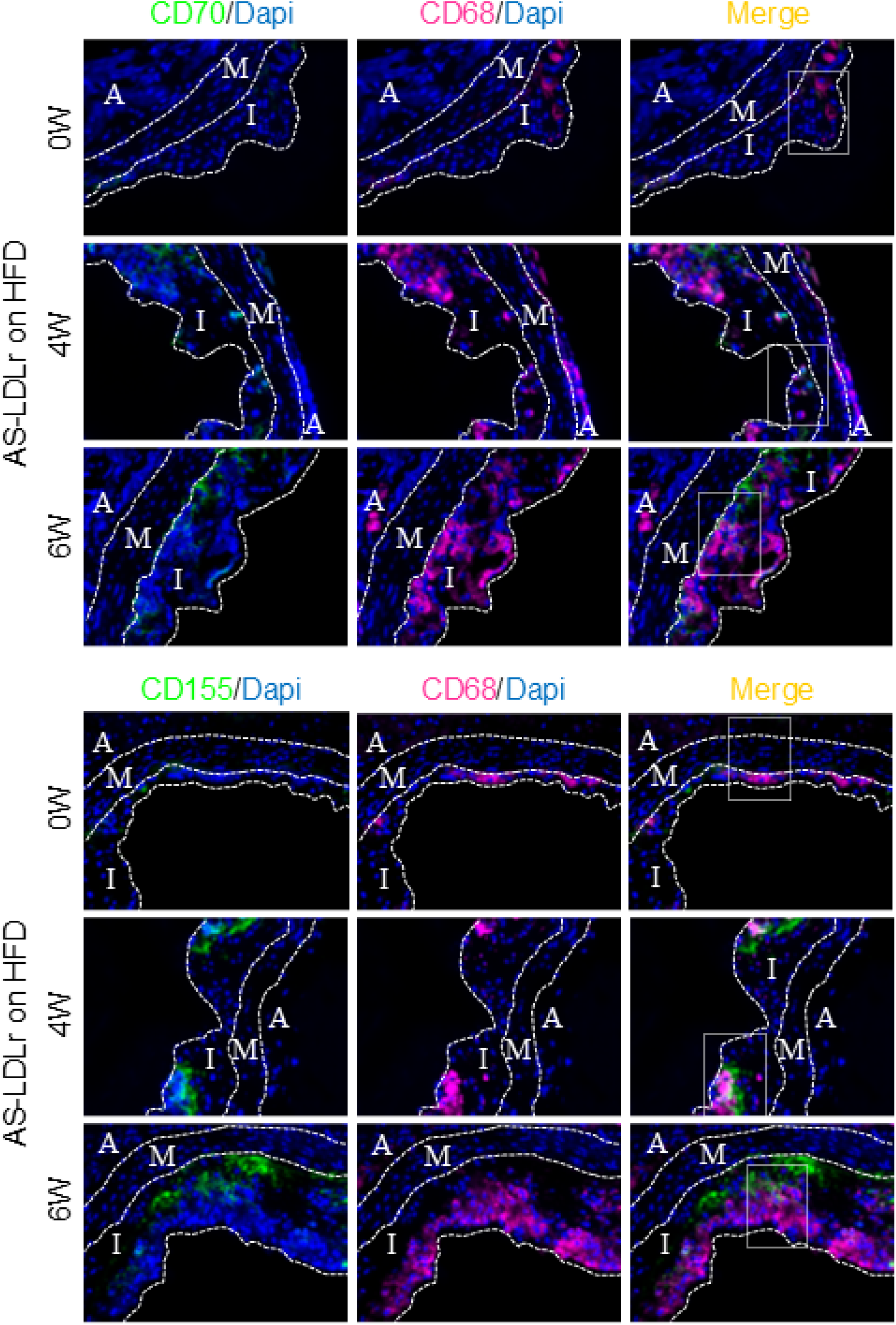
Lesional macrophages express CD70 and CD155 during early stages of atherosclerosis. Stressed macrophages are known to upregulate CD70 and CD155, while RAE-1 expression is predominantly restricted to non-haematopoietic cells, and lesion IgG is typically produced at distant sites. Immunofluorescence staining of atherosclerotic plaques collected from mice fed a high-fat diet for 0, 4, or 6 weeks revealed that lesional macrophages expressed CD70 and CD155 as early as week 4. Representative immunofluorescence images are shown; *n* = 5–8 mice per group. I: Tunica intima; M: Tunica media; A: Tunica adventitia.

**Supplementary Figure S3.**
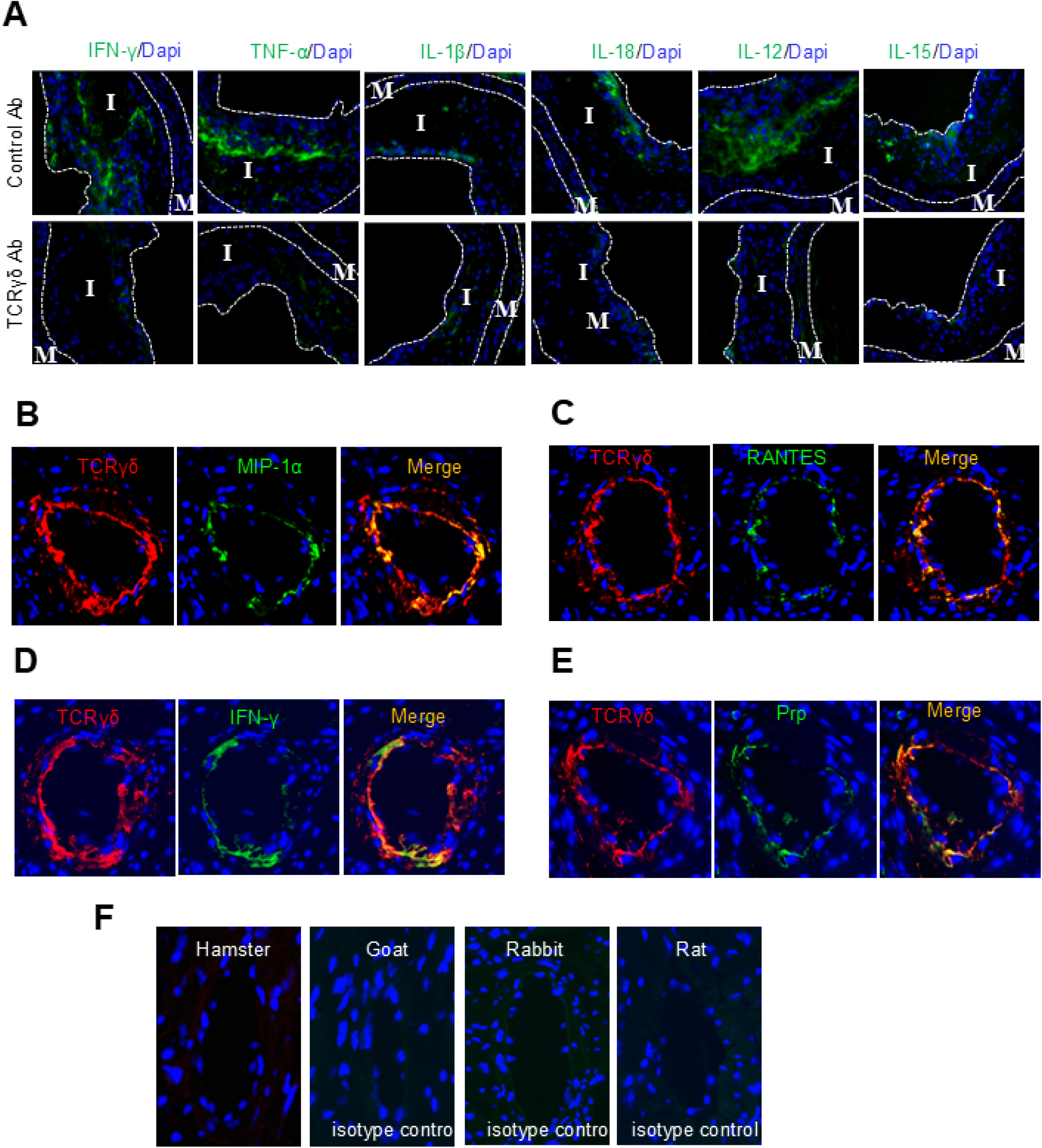
Coronary plaques contain γδ T cells expressing MIP-1α, RANTES, IFN-γ, and perforin. Proinflammatory cytokines, including IFN-γ, TNF-α, IL-1β, IL-18, IL-12, and IL-15, were reduced in atherosclerotic lesions following treatment with γδ T cell–depleting antibodies (**A**). To assess their mechanistic and therapeutic relevance in myocardial infarction, coronary plaques were collected from hyperlipidemic atherosclerotic mice with hypertension induced by transverse aortic constriction. Immunofluorescence staining was performed to evaluate γδ T cell colocalization with key effector molecules. Fluorescent imaging revealed that γδ T cells within coronary plaques expressed MIP-1α, RANTES, IFN-γ, and perforin (**B– E**). Control staining with isotype-matched antibodies is shown (**F**). Representative immunofluorescence images are presented; *n* = 5–8 mice per group. I: Tunica intima; M: Tunica media; A: Tunica adventitia.

**Supplementary Figure S4.**
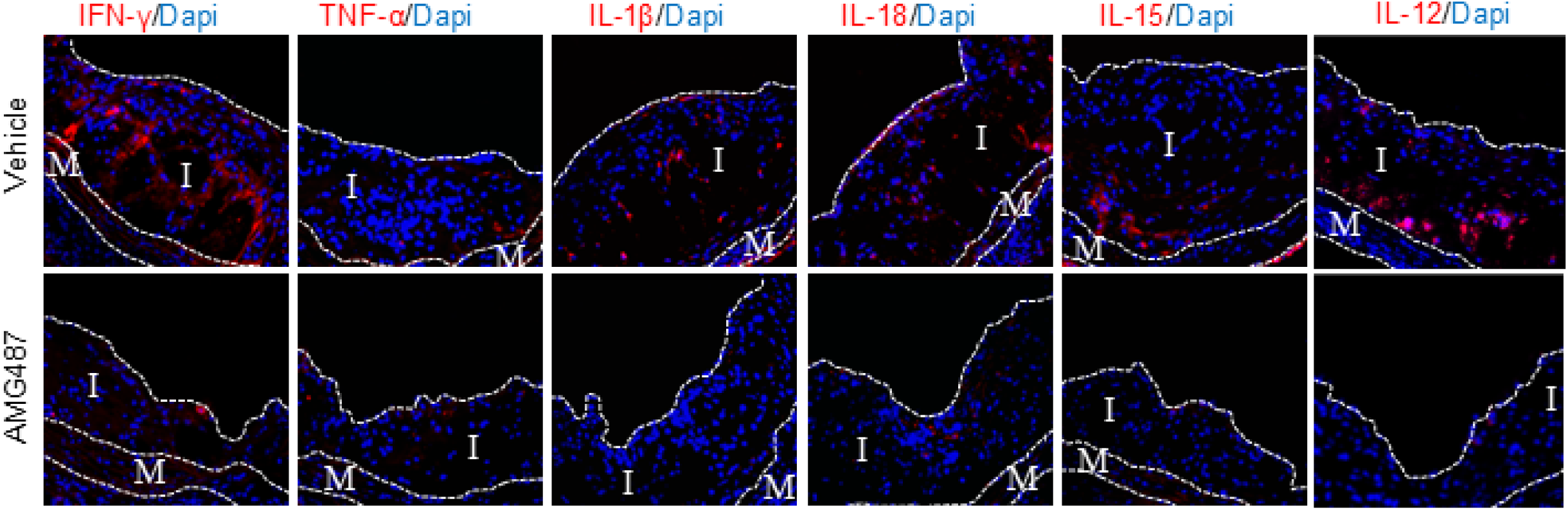
AMG487 treatment reduces plaque inflammation. Immunofluorescence staining of atherosclerotic lesions demonstrated reduced lesion inflammation as assessed by pproinflammatory cytokines, including IFN-γ, TNF-α, IL-1β, IL-18, IL-12, and IL-15. Representative immunofluorescence images are shown; *n* = 5–8 mice per group. I: Tunica intima; M: Tunica media; A: Tunica adventitia.

